# ImmuneRegulation: A web-based tool for identifying human immune regulatory elements

**DOI:** 10.1101/468124

**Authors:** Selim Kalayci, Myvizhi Esai Selvan, Irene Ramos, Chris Cotsapas, Ruth R. Montgomery, Gregory Poland, Bali Pulendran, John Tsang, Robert J. Klein, Zeynep H. Gümüş

## Abstract

Humans can vary considerably in their healthy immune phenotypes and in their immune responses to various stimuli. We have developed an interactive web-based tool, ImmuneRegulation, to enable discovery of human regulatory elements that drive some of the phenotypic differences observed in gene expression profiles. ImmuneRegulation currently provides the largest centrally integrated resource available in the literature on transcriptome regulation in whole blood and blood cell types, including genotype data from 23,040 individuals, with associated gene expression data from 30,562 experiments, that provide genetic variant-gene expression associations on ∼200 million eQTLs. In addition, it includes 14 million transcription factor (TF) binding region hits extracted from 1945 TF ChIP-seq peaks and the latest GWAS catalog of 67,230 published SNP-trait associations. Users can interactively explore ImmuneRegulation to visualize and discover associations between their gene(s) of interest and their regulators (genetic variants or transcription factors) across multiple cohorts and studies. These regulators can explain cohort or cell type dependent gene expression variations and may be critical in selecting the ideal cohorts or cell types for follow-up studies. Overall, ImmuneRegulation aims to contribute to our understanding of the effects of eQTLs and TFs on heterogeneous transcriptional responses reported in studies on the blood; in the development of molecular signatures of immune response; and facilitate their future application in patient management. ImmuneRegulation is freely available at http://icahn.mssm.edu/immuneregulation.

## INTRODUCTION

Recent high-throughput studies are contributing to an improved understanding of immune cell function and regulation (1). Extensive datasets are now available through the Human Immunology Project Consortium (HIPC) Phase I, which include measurements of healthy and activated human immune system, coupled with detailed clinical phenotyping in well-characterized cohorts (2–5). HIPC Phase 2 is continuing to collect data. In addition, large collaborative studies, including Genotype-Tissue Expression Project (GTEx) (6), Framingham Heart Study (7) and Encyclopedia of DNA Elements (ENCODE) (8–10) are collectively generating thousands of sequencing-based, genome-wide measurements of the transcriptome, transcription regulatory regions, transcription factor binding, and others that define states of the genome in many cell types and tissues, including a significant number of immune cell types. Furthermore, GWAS catalogs (11) provide a collection of large-scale studies that yield associations between genetic variants and various traits.

The wealth of data generated by these studies can be utilized for a deeper understanding of the mechanisms by which gene expression is regulated in different immune cells and immune systems of individuals in steady state and in response to immune stimuli (e.g. pathogens; disease; vaccines). Genetic variants typically appear to localize in the regulatory regions of genes and alter gene expression levels both proximally (putatively cis-acting) and distally (putatively trans-acting) (12), while transcription factors tend to bind directly to promoter regions proximally upstream of a gene, or directly to the RNA polymerase molecule and alter gene expression (13). However, often in the discovery phase, it is difficult to assess the real biological role of these regulatory elements without extensive experimentation on a large number of humans or immune cell types. To take advantage of the already generated rich datasets in the context of metadata that accompany them, a user-friendly immune-specific platform is needed to integrate, visualize and interactively explore these datasets. Such a platform can support investigations on human immunity, infection and disease as well as in the development of predictive models and therapies.

We have built an interactive web-based visual interface, ImmuneRegulation that capitalizes on the existing knowledge on the associations between transcription regulatory elements and gene expression changes in well-defined cohorts. ImmuneRegulation enables the exploration of massive immune-system specific gene regulation datasets and provides a centralized repository that enables seamless integration of information gained from large publicly available dataset resources to help efforts in understanding the associations between the immune system and its regulation. Easy visual queries to interact with these datasets assist discovery by creating resources for hypothesis generation. Users can simultaneously query multiple resources to identify cell-type or cohort specific regulatory elements that drive the expression of genes or gene sets of interest; study potential inherited susceptibility to specific responses or disease; or design follow-up cohort response studies with improved sensitivity and accuracy of classification by excluding or including certain individuals with specific genetic variants.

ImmuneRegulation currently includes genotype data from 23,040 individuals, associated with gene expression data from 30,562 experiments, providing genetic variant- gene expression associations on ∼200 million eQTLs. These enable the identification of gene(s) whose expressions are affected by human germline genetic variation in blood or blood cell types. Additionally, Transcription Factors (TFs) from publicly available ENCODE Consortium TF ChIP-seq studies in human are included. immuneRegulation currently incorporates around 14 million gene region hits extracted from 1,945 TF ChIP-seq peaks datasets. Furthermore, GWAS catalog data within immuneRegulation repository is comprised of 67,230 published SNP-trait association information.

In summary, ImmuneRegulation is a unique resource focused on regulation of gene(s) that are phenotype-dependent in expression response to immune stimuli. More specifically, it is an interactive web-based visual tool that enables the exploration of massive data on the regulation of immune-system specific genes in well-defined cohorts. Furthermore, immuneRegulation is currently the only visual interface for exploration of trans-eQTLs. Prior to ImmuneRegulation, these datasets were publicly available in separate resources without any exploratory features. Overall, ImmuneRegulation provides a centralized repository that enables seamless integration of information gained from HIPC with public dataset resources to help efforts in understanding the associations between the immune system and its regulation, where users can capture regulatory elements that drive some of the phenotypic differences observed in immune-related transcriptomes. ImmuneRegulation is available at http://icahn.mssm.edu/immuneregulation.

## MATERIAL AND METHODS

immuneRegulation provides a unified platform for a massive collection of immune regulation data in multiple cell types and in large human cohorts. Its visually intuitive and user-friendly interface helps users query, browse and interactively explore gene regulation datasets. Users can query for regulatory elements of genes of interest in real-time, utilizing its massive dataset. We illustrate the data repository and implementation details in the following sections.

### Data Collection and Processing

#### Publicly available eQTL datasets

We collected and curated multiple studies on blood or blood cell types from public datasets that are already available in literature. The complete list of these public datasets is provided in Table 1.

**Table 1.**
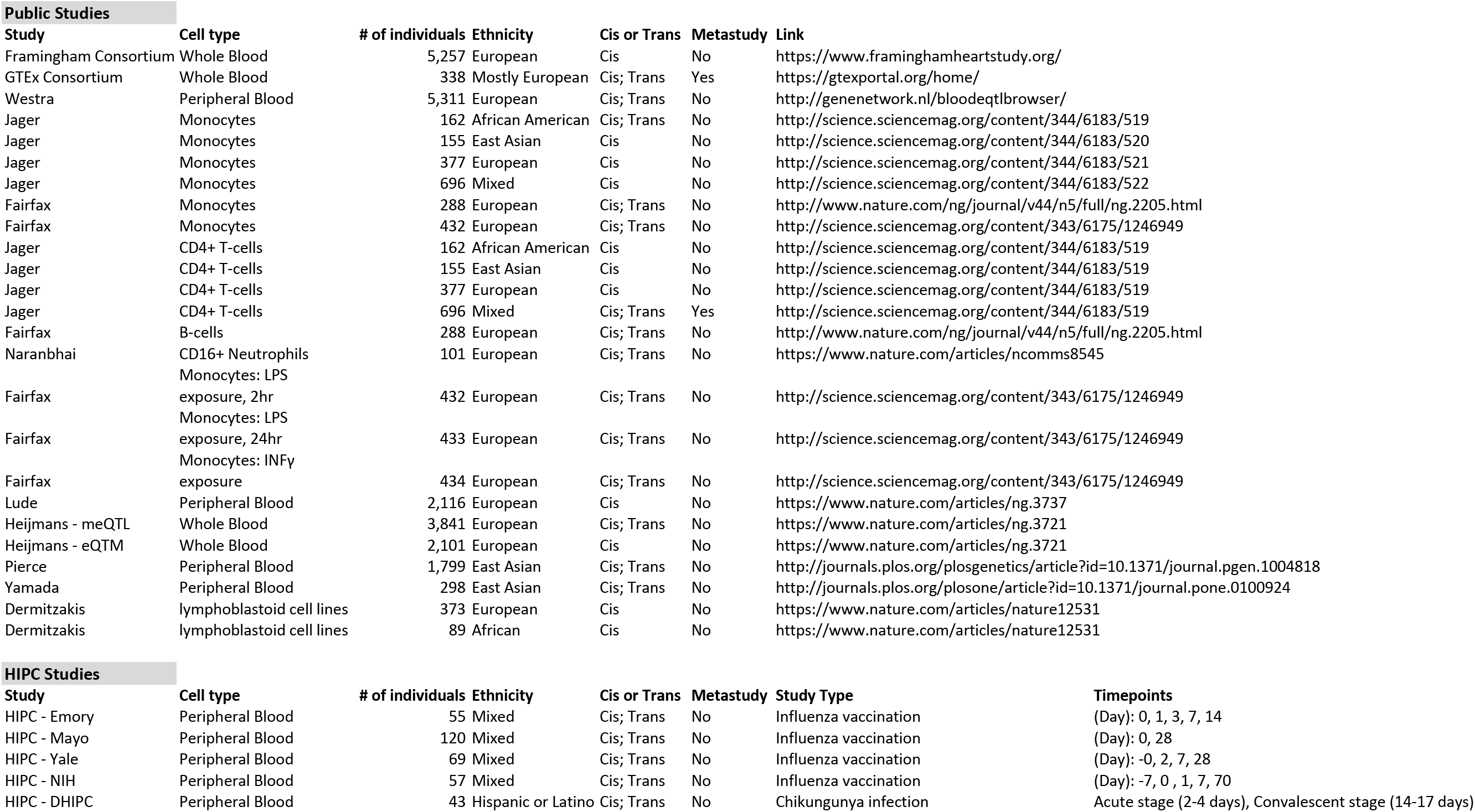
List of eQTL datasets provided in immuneRegulation.

#### cis- and trans-eQTLs from HIPC genotype-gene expression datasets

For HIPC studies that included both genotype and gene expression data (either microarray or RNA-seq), we first identified eQTLs by utilizing the pipeline provided at https://github.com/molgenis/systemsgenetics/wiki/eQTL-mapping-analysis-cookbook and then added them to the immuneRegulation database. These studies currently include systemic profiling of influenza-vaccinated individuals at multiple timepoints (Table 1).

#### cis- and trans-eQTLs from HIPC RNA-seq datasets

For HIPC RNA-seq studies without accompanying genotype data, we downloaded all raw datasets from GEO (www.ncbi.nlm.nih.gov/geo) aligned spliced reads to human genome via STAR (14), and performed SNP and indel calling by using the Genome Analysis Toolkit (15) (GATK, https://www.broadinstitute.org/gatk/) best practices workflow, before running the same eQTL analysis pipeline. These studies currently include datasets from Chikungunya virus infected individuals profiled at both acute and convalescent stages of the disease (Table 1).

#### Transcription Factors (TF) datasets

We have used all human TF datasets from the publicly available ENCODE Consortium ChIP-seq studies (16, 17). These include a total of 1,945 ChIP-seq TF-binding site datasets (in bed narrowPeak file format) from 144 different cell/tissue types and 736 different TF targets. For each dataset, we calculated gene region hits using bedtools v2.25.0 (18). Then, for each gene, we incorporated the metadata of all datasets with gene region hit(s). The specific data tracks are transferred from ENCODE portal (16) upon user request.

#### GWAS datasets

We have incorporated the complete NHGRI-EBI GWAS catalog (11) (www.ebi.ac.uk/gwas, downloaded in March 2018) into ImmuneRegulation data repository to be utilized in queries. Currently, this includes information about 67,230 published SNP-trait associations.

### Implementation and Interface

#### Implementation

ImmuneRegulation utilizes multiple client-side Javascript libraries (e.g. D3.js, JQuery, etc.) to visualize and interact with large volumes of multiple data types in real time. For visualizations that are displayed in the genome browser, we incorporated igv.js (https://igv.org/doc/doc.html) and customized it to serve our specific needs. We also incorporate several other utility libraries (e.g. dataTables.js, underscore.js, etc.) for data manipulation and interaction. For styling of the web interface, we mainly relied on Bootstrap v3.3.7 and our own custom CSS elements. As a result, immuneRegulation runs on all modern web browsers.

#### Interface

We designed the immuneRegulation interface with the aim of guiding users during queries and interactions with results. Warning messages are displayed dynamically to help users construct the queries properly. Similarly, certain dynamic elements are displayed as the users interact with the page and help inform the users regarding any possible interactivity option or further information. We also incorporated multiple example queries into immuneRegulation interface for users to gain familiarity with the tool and to create customized queries of their own easily. Furthermore, multiple detailed tutorials help explain various components and functionalities within immuneRegulation. An FAQ page provides basic information pertaining to the content as well as technical aspects of the interface (e.g. missing data, load time, etc.). Users can provide feedback or ask questions regarding any technical or scientific matters directly from the immuneRegulation interface as well.

## RESULTS

ImmuneRegulation centrally integrates and utilizes massive immune-specific datasets in the context of their rich metadata. In its front-end, a visually intuitive web interface enables query, browsing and interaction with large volumes of data. Users can query for regulatory elements for gene(s) of interest in real-time. For gene(s) queried, visual, interactive summaries of regulatory elements are returned to help explore, contextualize and communicate statistical analyses and results.

### Data Repository

Since expression regulation is tissue-specific, we provide an immune system focused resource to understand gene regulation in whole blood or blood cells (T-cells, macrophages and monocytes). ImmuneRegulation currently includes genotype data from 23,040 individuals, associated with gene expression data from 30,562 experiments, providing genetic variant- gene expression associations on ∼200 million eQTLs. These enable the identification of gene(s) whose expressions are affected by human germline genetic variation in blood or blood cell types. Additionally, Transcription Factors (TFs) from publicly available ENCODE Consortium TF ChIP-seq studies in human are included. immuneRegulation currently incorporates around 14 million gene region hits extracted from 1,945 TF ChIP-seq peaks datasets. Furthermore, GWAS catalog data within immuneRegulation repository is comprised of 67,230 published SNP-trait association information.

### Getting Started (Performing Queries)

The landing page of immuneRegulation provides the interface to construct queries as displayed in Figure 1. Users can enter gene(s) of interest by providing the associated HUGO gene symbol(s) within the text-box located at the top of the interface. (In this version, users are allowed to enter multiple genes for trans-eQTL queries; whereas only one gene input is allowed for cis-eQTL or TF queries) Then, users can select study type on the left (HIPC: cis-eQTL or trans-eQTL; or non-HIPC: cis-eQTL or trans-eQTL; or Transcription Factors) and filter relevant datasets among the collection of datasets listed on the right. Users can look through the list of datasets by referring to basic information (sample size; summary; link to publication or study) provided adjacent to each dataset. After selecting one or more datasets, user can click the Submit Query button. For each study type (cis-eQTL; trans-eQTL; Transcription Factors), users will get different visuals to display the results for their queries, which are explained below. Users can perform an updated or a completely different query by clicking the Modify Query button and making the desired changes.

**Figure 1.**
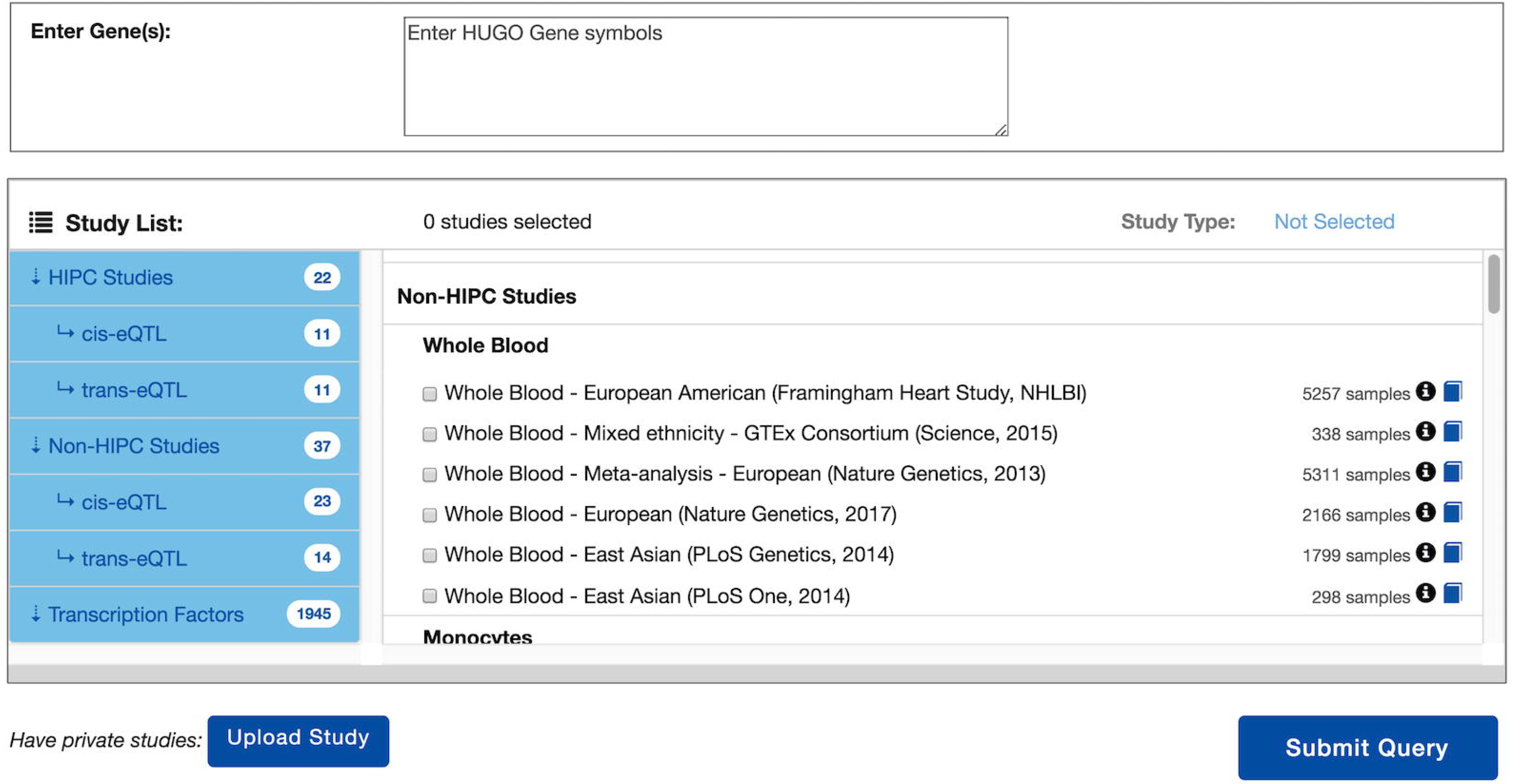
Main visual interface for performing queries.

### Querying cis-eQTL Data

Users can enter a single gene and select one or more relevant datasets to perform a cis-eQTL query. This opens our customized Integrative Genomics Viewer (IGV) browser (19) as shown in Figure 2; cis-eQTLs for each selected dataset are displayed in separate genome browser tracks with their associated p-values. Red circles indicate that the cis-eQTL appears in multiple tracks; blue circles indicate that the cis-eQTL appears only in one track; and grey circles represent cis-eQTLs for a different gene. User can click on any specific cis-eQTL and a pop-up dialog box displays more details (position, gene, p-value, SNP ID and information link) about it. Users can zoom in and out, and also move across the genomic region to navigate other information. Additionally, two more tracks are displayed below cis-eQTL tracks for further information and reference. GWAS Catalog track displays SNPs associated with any GWAS findings in the genomic region. Clicking on any specific GWAS finding opens a pop-up dialog box displaying further details, such as disease/trait and publication link, regarding the GWAS entry. Genes track displays gene isoforms and provides a quick view of the relative locations of other information displayed in the genome browser.

**Figure 2.**
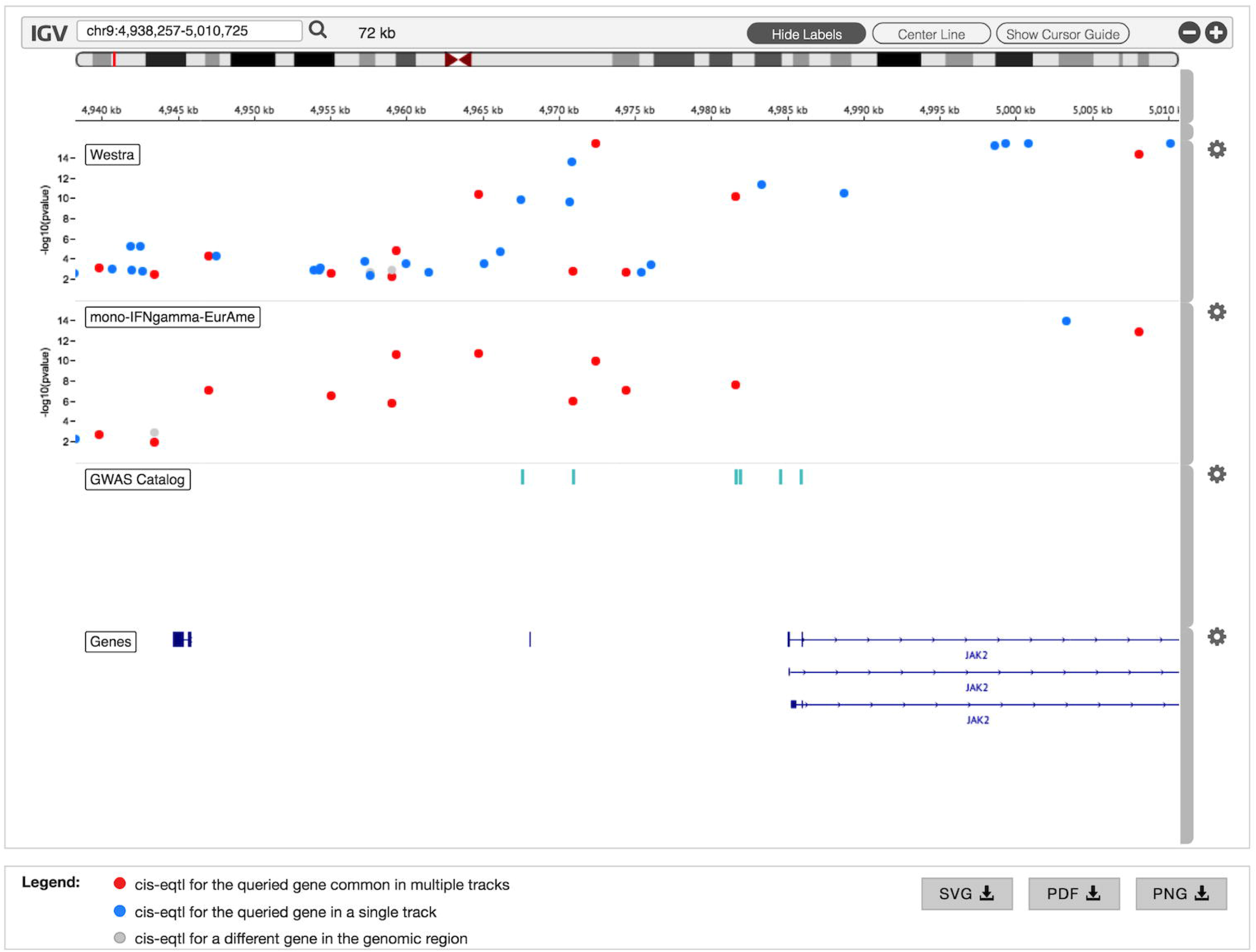
cis-eQTL results for two different datasets displayed within the customized IGV browser. GWAS Catalog and Genes tracks are also displayed by default.

### Querying *trans-eQTL* Data

Users can enter gene(s) of interest and select one or more relevant datasets to perform a trans-eQTL query. If multiple genes are provided as input, user is given the option, via a dynamical visual element, to apply either intersection or union operation on the gene list. In case where the user selects the intersection operation, only those SNPs that act as trans-eQTLs for more than one gene are displayed as a table. However, if the union operation is selected for multiple genes or if a single gene is provided as input, results are visualized similar to Figure 3. Figure 3 Panel A displays a screenshot of the resulting interactive graph of all trans-eQTLs that regulate the query genes, rank-ordered and visualized based on their p-values; users can also view their genomic locations in a Manhattan plot (Panel B). Users can filter trans-eQTL results based on genomic locations of SNPs, as well as using a subset of genes. Black circles indicate trans-eQTLs that have associated GWAS hits; blue circles indicate trans-eQTLs without any GWAS hit associations. Hovering over any specific trans-eQTL finding provides a quick look into its details via a tooltip text. For deeper analyses, users can click on a specific trans-eQTL from either graph; a text-box as in Panel C appears to the right of the graph with details about the trans-eQTL and links to further associated information. Clicking on the first link with the down arrow opens up a table, as in Panel D, that displays other gene targets for the specific SNP associated with the trans-eQTL. Similarly, clicking on the second link with the down arrow opens up a table, as in Panel E, that displays GWAS hits associated with the specific SNP.

**Figure 3.**
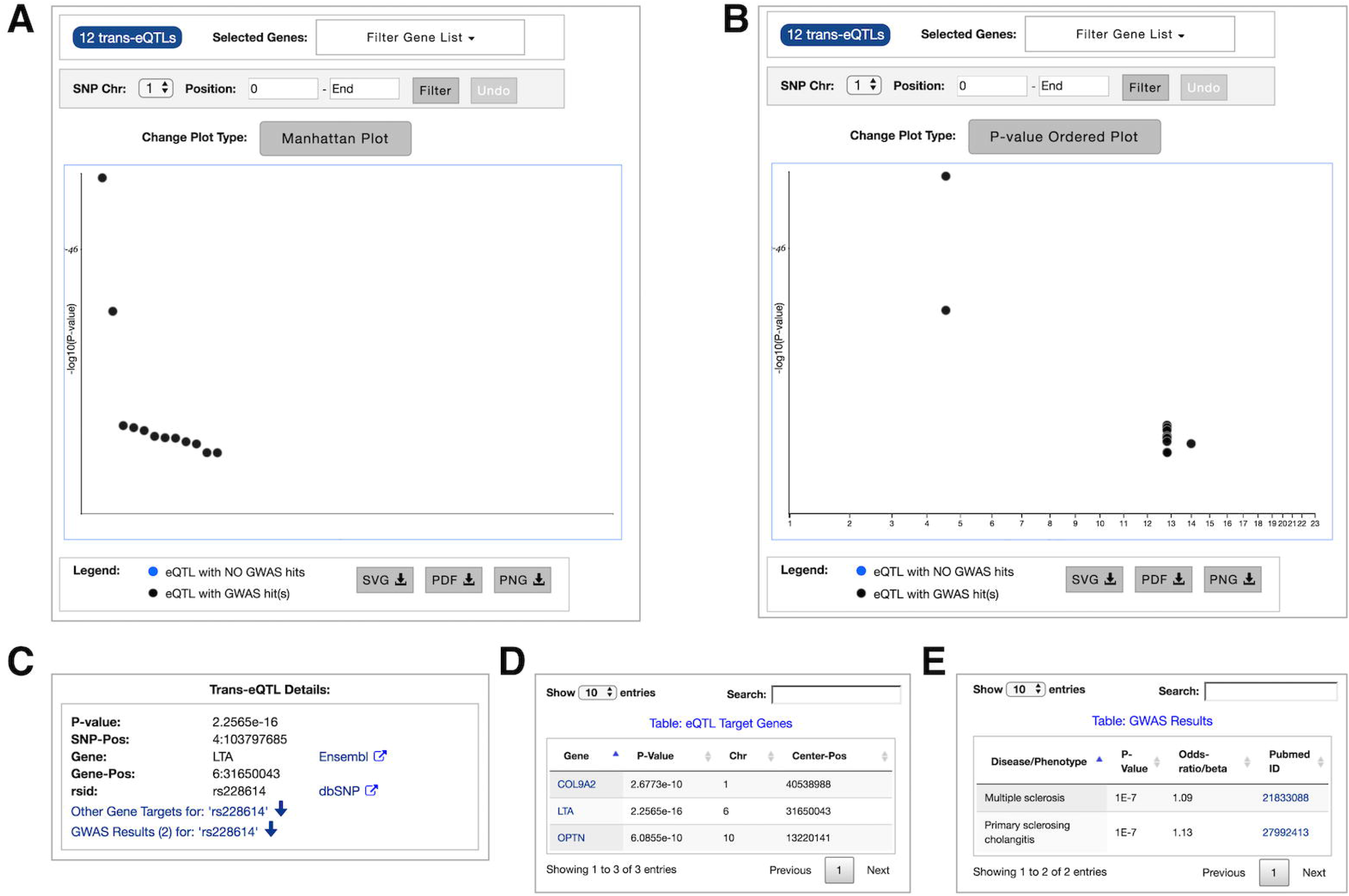
All trans-eQTL results from query **A** p-value ordered; **B** Manhattan plot; For a specific trans-eQTL **C** details table; **D** eQTL target genes table; **E** GWAS results table.

### Querying Transcription-Factor (TF) Data

Users can enter a single gene and select the ENCODE collection of datasets to query TFs. This opens a table, Figure 4 Panel A, that displays all datasets that have gene region hits for the gene queried. Each row in the table corresponds to a cell type-TF target combination and it can be filtered to look for a specific cell type or TF target using the Search box. Users can explore and select the datasets of interest as shown in Panel B. Clicking the Submit button in Panel B opens our customized IGV browser as shown in Panel C. TF ChIP-seq peaks for each selected dataset are displayed in separate genome browser tracks. GWAS Catalog and Genes tracks are also displayed to provide further information about the genomic region.

**Figure 4.**
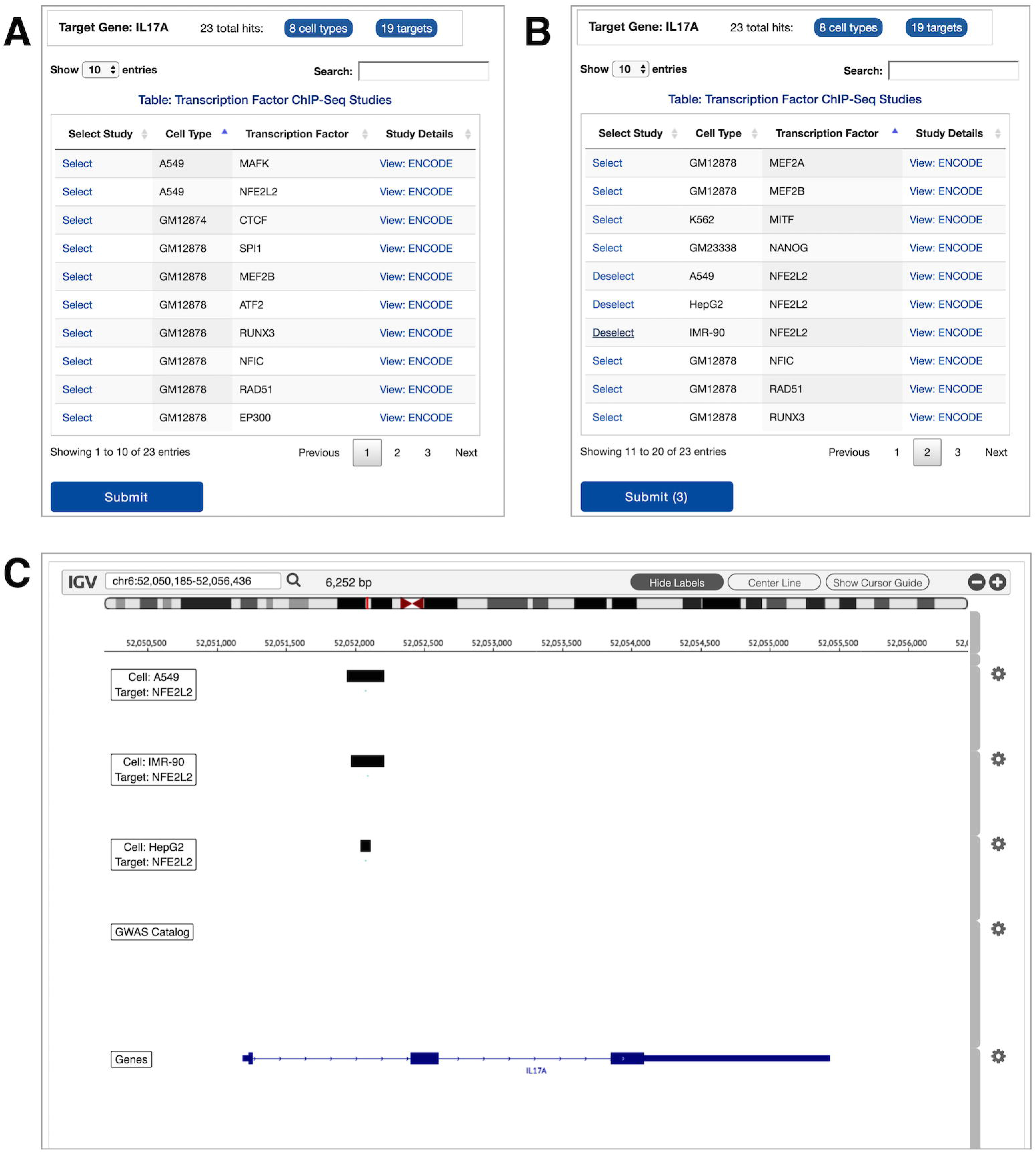
Transcription Factor gene hits results **A** table sorted by cell type; **B** table sorted by target, and visualization list is populated; **C** studies added to the list visualized within the customized IGV browser.

### Uploading and Visualizing Private Data

immuneRegulation provides capability for users to upload datasets of their own (cis-eQTL and trans-eQTL) and integrate with the rest of the interface. The important detail here is that the user datasets are not transferred to any remote server or third-party site; the operation is handled completely within the local browser interface of the user performing the upload. Users can utilize this functionality by clicking the Upload Study button in the main query browser interface. A pop-up window opens up with descriptions of the proper format for datasets that can be uploaded, and enables the users to upload one or more datasets (cis-eQTL and/or trans-eQTL). After the user saves the changes, those datasets are added to the query browser interface similar to as shown in Figure 5. These datasets are completely integrated within immuneRegulation interface; users can browse and query them in combination with any existing datasets, and interact with the visual results in the same way.

**Figure 5.**
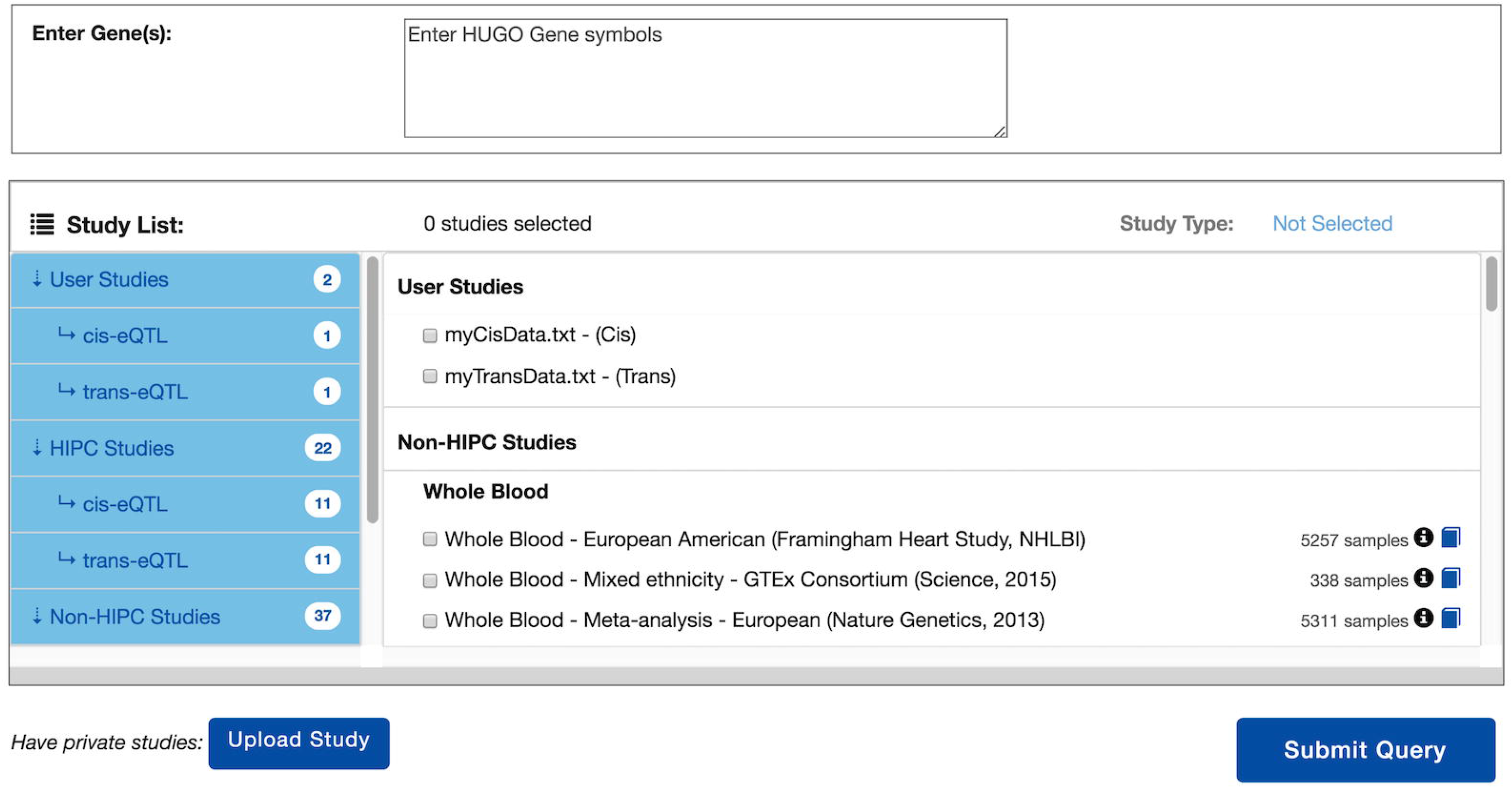
Query interface displaying user uploaded datasets.

## DISCUSSION

To comprehensively understand heterogeneity in the healthy human immune system and in response to stimuli, multiple groups are generating high-throughput measurements of gene regulation in the human immune system. These massive data create their own challenges in accurately interpreting immunophenotyping data, necessitating easy to use data mining and interpretation tools. An important goal in these efforts is phenotyping the immune states and diseases for diagnosis, prognosis and selection of therapies. ImmuneRegulation helps investigators interpret multiple data resources in their efforts to generate hypotheses on how transcriptome differences observed in different members of the same cohort can be explained by specific regulators. While such regulators (genetic variants or transcription factors) can yield insights into the immune response to stimuli, and can improve sensitivity and accuracy of classification, often in the discovery phase it is difficult to assess their real biological role without extensive experimentation. Integrating multiple datasets with existing genotype and transcriptome data on a large number of humans and immune cell types in a single, user-friendly interface supports investigations on human immunity, infection and disease. The availability of reliably curated patient cohorts and the integrative interface of ImmuneRegulation that provides exploration of massive immune regulatory datasets can help accelerate our understanding of the human immune system. We anticipate that rapid, intuitive and integrated analyses within our interface will significantly empower immune researchers in their study of the complex role genomic regulation plays in immune phenotype responses and help the translation of the findings to insights and applications in basic and clinical immunology. Specifically, some of the differences observed in the phenotypic responses may be caused by inter-individual genetic variation, and identifying the regulatory elements that lead to differences in transcriptome responses within these cohorts can yield insights into the development, remission and exacerbation of disease; they may as well improve diagnostic sensitivity and accuracy of cohort classification, and ultimately guide treatment.

## AVAILABILITY

ImmuneRegulation is an open source collaborative initiative available in the GitHub repository https://github.com/gumuslab-mssm/immuneRegulation

## ACKNOWLEDGEMENT

We thank Ana Fernandez-Sesma, Eun-Young Kim, Andrew Kasarskis, Steven Kleinstein, Chris Cotsapas, and Steven Wolinksy for their valuable feedback in helping us to better understand and address user needs in interacting with the tool.

